# Growth inhibition of *Akkermansia muciniphila* by a secreted pathobiont sialidase

**DOI:** 10.1101/2022.03.15.484512

**Authors:** Guus H. van Muijlwijk, Emma Bröring, Guido van Mierlo, Pascal W.T.C. Jansen, Michiel Vermeulen, Piet C. Aerts, Jos P.M. van Putten, Noah W. Palm, Marcel R. de Zoete

**Author notes:** Corresponding author: Marcel R. de Zoete.

## Abstract

*Akkermansia muciniphila* is considered a key constituent of a healthy gut microbiota. In inflammatory bowel disease (IBD), *A. muciniphila* has a reduced abundance while other, putative pathogenic, mucus colonizers bloom. We hypothesized that interbacterial competition may contribute to this observation. By screening the supernatants of a panel of enteric bacteria, we discovered that a previously uncharacterized *Allobaculum* species potently inhibits the growth of *A. muciniphila*. Mass spectrometry analysis identified a secreted *Allobaculum* sialidase as inhibitor of *A. muciniphila* growth. The sialidase targets sialic acids on casein *O*-glycans, thereby altering the accessibility of nutrients critical for *A. muciniphila*. The altered glycometabolic niche results in distorted *A. muciniphila* cell division and efficiently arrests its growth. The identification of a novel mechanism of *A. muciniphila* growth inhibition by a competing bacterial pathobiont may provide a rationale for interventions aimed at restoring and maintaining a healthy microbiota symbiosis in patients with intestinal disease.

## Introduction

The human intestinal tract is a highly dynamic ecosystem that is home to trillions of microbes, known as the gut microbiota^1^. Throughout our lives, the gut ecosystem is subject to various disturbing influences driven by external factors like changes in diet, exposure to antibiotics and competition with newly acquired commensals or invading enteropathogens^2–4^. All these events are reflected in the composition of the intestinal microbiota, and changes in this composition have been associated with numerous diseases^5–10^.

Intestinal bacteria can significantly affect host physiology, dependent on their metabolism and lifestyle^11–13^. Due to their position in the gut ecosystem, mucus-colonizing and mucosa-associated bacteria are more likely to affect host immune responses and have a dominant effect on host physiology and the development of intestinal disease^14–18^. One of the species known to reside in the mucus layer and to affect host immune responses is *Akkermansia muciniphila*^19,20^. This mucin-degrading bacterium plays an important role in the maintenance of gut immune homeostasis as it has been reported to improve mucus layer thickness and epithelial barrier function^20^. Furthermore, it induces immune tolerance through the induction of Tregs and is often highly coated by secretory IgA, suggesting close interactions with the intestinal immune system.^18,20–23^ Notably, a reduced abundance of *A. muciniphila* has been reported in inflammatory bowel disease (IBD) patients and several other inflammatory diseases such as type 2 diabetes and metabolic syndrome, but the reasons for this reduced abundance remain unclear^20,24,25^. Identification of the factors that negatively impact the abundance of *A. muciniphila* may provide important new insights into the mucosal bacterial ecosystem and its effects on the host.

Multiple factors may contribute to the loss of *A. muciniphila*, such as changes in diet or the excessive use of antibiotics^26–29^. Another factor that is often overlooked, and which may act in synchrony with other factors, is the possibility of direct interbacterial antagonism by bacteria occupying the same niche. Given that many such interactions are known to exist in the bacterial kingdom, we investigated the possibility that distinct bacteria present in the gut of IBD patients have the ability to inhibit *A. muciniphila* directly. By screening the interaction of *A. muciniphila* with a panel of IBD-associated bacteria we discovered that a secreted protein from the pathobiont *Allobaculum mucolyticum* potently inhibits the growth of *A. muciniphila*. This protein was identified to be a sialidase that desialylates *O*-glycans on substrates required for *A. muciniphila* growth. The alteration of the glycometabolic niche efficiently arrests bacterial growth through distorted bacterial cell division. The identification of a novel mechanism of *A. muciniphila* growth inhibition by a competing bacterial pathobiont may pave the road for therapies aimed at restoring and maintaining microbiota homeostasis in patients with intestinal disease.

## Materials & Methods

### Bacterial culturing conditions

All bacteria were routinely cultured at 37°C under strict anaerobic conditions at an atmosphere of 5% H2, 10% CO_2_ and 85% N_2_. Unless stated otherwise, bacteria were cultured in enriched Gut Microbiota Medium (GMM) broth, of which the recipe is described in supplementary Table 1. The defined minimal medium (DMM) was derived from GMM and prepared using the following basal ingredients at similar concentrations as in GMM: Phosphate buffer, Vitamin K solution, TYG salts, CaCl_2_, FeSO_4_-7H_2_O, Resazurin, Histidin-Hematin solution, Trace Mineral supplement, Vitamin supplement and L-Cysteine HCl. This basal medium was supplemented with trypticase peptone (10 g/L) and glucose (10 mM). Prior to use for bacterial growth, all media were prereduced in the anaerobic chamber for at least 12 h, unless stated otherwise.

### Bacterial conditioned media (CM)

Bacterial conditioned media (CM) were collected from a panel of 14 different enteric bacterial isolates previously isolated from IBD patients^18^. For each strain, conditioned medium was collected from three biologically independent broth cultures started from a glycerol stock. Cultures were grown statically for 48 h after which the conditioned media were harvested by centrifugation, filter-sterilized using 0.2 μm filter and stored at −20°C prior to use.

### *A. muciniphila* growth inhibition experiments

*Akkermansia muciniphila* (ATCC BAA-835) and an independent *A. muciniphila* clinical isolate were routinely cultured at 37°C under anaerobic conditions as described above.

#### A. muciniphila starter cultures

All growth assays were preceded by growing *A. muciniphila* broth starter cultures. For this, 3 mL of prereduced GMM broth was inoculated with *A. muciniphila* from a glycerol stock. After 48 h of growth, bacteria were pelleted by centrifugation (3,000 x g, 5 min) and gently washed twice with the desired medium used in the growth assay.

For all the following experiments *A. muciniphila* was diluted in the desired medium and cultured in a 96-Well Multiwell Flat Bottom Plate (Corning Costar) with a final volume of 200 μL per well and a starting optical density (600 nm) of 0.01. After 48 h, or unless otherwise stated, the plate was removed from the anaerobic chamber and final absorbance values (600 nm) were recorded using a FLUOstar Omega plate reader (BMG Labtech). Values were corrected for background levels in respective negative controls.

#### A. muciniphila cultured with bacterial conditioned media

Sterile bacterial CM were divided into two fractions. One fraction was kept at room temperature (21°C), while the other fraction was heat-inactivated (98°C, 30 min), after which both fractions were transferred to the anaerobic chamber. 50 μL of the active and heat-inactivated fractions were transferred to a 96-Well Multiwell Plate (Corning Costar) per well in duplicates to which *A. muciniphila* starter cultures were added for final volume of 200 uL.

### Treatments and supplementation of *Allobaculum mucolyticum* conditioned medium

For clarity *Allobaculum mucolyticum* will hereafter solely be referred to as *Allobaculum*. All treatments were performed prior to addition of CM to *A. muciniphila* cultures, unless otherwise stated.

#### Proteinase K treatment

*Allobaculum* CM or GMM control media were treated with Proteinase K (1 mg/mL), PMSF (1 mM) or PMSF-inactivated Proteinase K (1 mM and 1 mg/mL, respectively) for 16 h at 37°C, after which the Proteinase K-treated samples were supplemented with PMSF (1 mM).

#### Size-fractionation

*Allobaculum* CM was fractionated based on size through successive use of 50, 30, and 3 kDa MWCO Amicon Ultra filters (Thermo Fisher Scientific). Intermediate retentates were reconstituted to the starting volume using GMM.

#### Size-fractionation by high-resolution chromatography

Twenty mL of *Allobaculum* CM was concentrated to 0.5 mL using a 50 kDa MWCO Amicon Ultra filter (Thermo Fisher Scientific), reconstituted to 20 mL using PBS and reconcentrated once more to 0.5 mL. High performance liquid chromatography (HPLC) of this fraction was conducted using a Superdex^®^ 200 Increase 10/300 GL on the ÄKTA pure protein purification system (GE Healthcare, Uppsala, Sweden). The proteins were eluted with PBS (pH 7.4) at a flow rate of 0.75 mL/min, detected by absorbance at 280 nm, and collected in 0.5 mL fractions. Fractions containing protein peaks (fractions 13-56) were tested for growth inhibition of *A. muciniphila*. For this, 100 μL of each size-exclusion fraction was combined with *A. muciniphila* diluted in 2x concentrated GMM and cultured as described above. The surplus of the size-exclusion fractions was stored at −20°C prior to further fractionation.

#### Fractionation by anion-exchange chromatography

Inhibiting size-exclusion fractions (fractions 17-28) were pooled to a total volume of 3 mL. Pooled fractions were dialyzed overnight at 4°C, against starting buffer A (20 mM Bis-Tris, pH 6) using a 10 kDa MWCO dialysis tube. High performance liquid chromatography (HPLC) was conducted using a HiTrap Q XL 1 mL Sepharose column on the ÄKTA pure protein purification system (GE Healthcare, Uppsala, Sweden). Proteins were eluted with starting buffer A followed by a gradually increasing concentration of buffer B (20 mM Bis-Tris, 1 M NaCl, pH 6) at at a flow rate of 0.75 mL/min, detected by absorbance at 280 nm, and collected in 0.5 mL fractions. Fractions or matching NaCl molarity controls in TBS pH 6 were tested for growth inhibition of *A. muciniphila*. For this, 50 μL of each anion-exchange fraction was diluted with 50 μL MilliQ and combined with *A. muciniphila* diluted in 2x concentrated GMM and cultured as described above. The surplus of the size-exclusion fractions was stored at −20°C prior to further processing.

#### Supplementation with defined monosaccharides

L-Fucose (Sigma), D+Galactose (Sigma), N-Acetylneuraminic acid (Carbosynth), N-acetyl-D-glucosamine (GlcNAc, Sigma), and N-acetyl-D-galactosamine (GalNAc, Carbosynth) were dissolved in MilliQ to a final concentration of 100 mM, adjusted to pH 7.2 using 1 M NaOH or HCl if necessary, and filter-sterilized using 0.2 μm filter. Zero, 25 or 50 μl of active or heat-inactivated *Allobaculum* CM or GMM control, for a final concentration of 0%, 12.5% and 25% supernatant, respectively, were combined with 50 μl of each monosaccharide solution, a dilution thereof, or MilliQ control for a final concentration of 0 mM, 1 mM, 5 mM or 25 mM monosaccharide. Mixtures were combined with *A. muciniphila* previously diluted in 2X concentrated GMM.

### NanH1 pretreatment of GMM and DMM

Recombinant NanH1 or heat-inactivated NanH1 (2.5 μg/mL) or an equal volume of PBS (mock) as control were added to GMM or DMM. Mixtures were incubated for 16 h at 37°C after which they were heat-inactivated (98°C, 30 min) prior to use.

### *A. muciniphila* growth with recombinant *Allobaculum* NanH1 sialidase and other bacterial sialidases

Recombinant NanH1, heat-inactivated NanH1 (98°C, 30 min) and the mutant NanH1ΔRIP (see below) were diluted to desired concentration in 100 μL MilliQ or MilliQ supplemented with sialic acid (Neu5Ac) and combined with *A. muciniphila* in 2x concentrated GMM. Recombinant *Allobaculum* NanH2, CPNA (Sigma), VCNA (Roche), and AUNA (Roche) were also diluted in MilliQ and the concentration was adjusted to equal the sialidase activity of NanH1 on the 4-MU-NANA substrate (see below).

### Sialidase activity assays

Fifty μL of *Allobaculum* CM or GMM control were added to a 96-Well Multiwell flat bottom plate (Corning Costar). Recombinant *Allobaculum* NanH1, NanH1ΔRIP, NanH2 (all at 500 ug/mL), and CPNA (25 U/mL), VCNA (16 U/mL) and AUNA (10 U/mL) were diluted to a range of different concentrations in GMM after which 50 μL was added to the wells. Next, 50 μL of 4-MU-Neu5Ac substrate (200 μM in MilliQ, Cayman) was added to each well after which the plate was transferred to the FLUOstar Omega plate reader (BMG labtech), which was kept at 37°C. Fluorescence was recorded over time or as an endpoint measurement (indicated in graph) using the 340 ex / 460 em filter set. Relative sialidase activities in GMM for stock solutions of NanH1 / NanH1ΔRIP / NanH2 / CPNA / VCNA / AUNA are 100% / 0% / 8.4% / 78.5% / 118.9% / 99.5%, respectively, relative to NanH1.

### Mass spectrometric analysis of fractionated *Allobaculum* conditioned media

#### In-gel digestion

The strongest inhibiting anion-exhange fraction (fraction 50), and a non-inhibiting fraction (fraction 12) were analysed using LC-MS/MS. Proteins mixtures were denatured in 1X Nu-PAGE LDS sample buffer (NP0007, Invitrogen) and separated using a SDS PAGE gel. Gels were fixed with 10% acidic acid and 50% methanol, and stained with the Colloidial Blue Staining Kit (Invitrogen), followed by staining with colloidal Blue (20% stainer A (Invitrogen 46-7015) and 5% stainer B (Invitrogen 46-7016) in the presence of 20% methanol. Gels were destained with an excess of deionized water for 30 min at RT. Bands of interest were excised and at the same time control pieces of unstained gel fractions were taken along as controls. Each slice was transferred to an 1.5 ml Eppendorf tube and further destained in a mixture of 25 mM ammonium bicarbonate (ABC) and 50% ethanol. Slices were then washed with 1 mL Acetonitril (ACN) for 10 min at, 1 mL ABC (50 mM) and again twice with ACN. All liquid was removed and tubes were air dried using a speedvac for 10 min. Then 200 μL of 10 mM dithiothreitol (DTT) in ABC was added and tubes were incubated for 45 min at 50 °C. This solution was removed and the gel was resuspended in 300 uL of 50 mM IAA in ABC followed by a 30 min incubation in the dark. After a wash with ABC and two washes with ACN, tubes were air dried once more using a speedvac and gel slices were coved in a Trypsin / ABC mixture followed by overnight digestion. Next, digested peptides were extracted using two 15 min incubations in a 100 μL mixture of 30% ACN and 3% formic acid in water, and two 15 min incubations in 100 μL of ACN. Resulting supernatants were combined (yielding ~500 μL) and concentrated to 100 μL using a speedvac. The resulting peptide was desalted using StageTips^30^.

#### LC-MS/MS measurements and data analysis

Digested peptides were analysed using an Easy-nLC1000 (Thermo) connected to a LTQ-Orbitrap-Fusion (Thermo). Raw files were analysed using standard settings of MaxQuant 1.5.1.0. Options LFQ, iBAQ and match between runs were selected. Perseus 1.5.1.0.15 was used to filter out proteins flagged as contaminant, reverse or only identified by site.

### Fluorescence confocal microscopy

*A. muciniphila* was grown for 48 h in GMM with 25% active or heat-inactivated *Allobaculum* conditioned medium or in either mock or NanH1 pretreated GMM, as described above. Bacteria were pelleted (3,000 x *g*, 5 min, at room temperature (RT)), washed once in PBS pH 7.4, resuspended in PBS + 4% paraformaldehyde (PFA) and fixed for 15 min at RT. Bacteria were washed as described above, twice in PBS, once in MilliQ water, and stained for 10 min at RT with 5 μM SYTO-9 dye (ThermoFisher) in MilliQ, while shielded from light. Five μL of each sample was spread on a poly-L-lysine-coated coverslip, air dried for 20-30 min at RT, mounted on a microscope slide using ProLong Diamond Antifade Mountant (Invitrogen) and allowed to dry overnight, while shielded from light. Samples were imaged on a Leica SPE-II laser confocal microscope using 63.0× 1.40 objective, and images were processed using Leica LAS AF software.

### Scanning electron Microscopy (SEM)

*A. muciniphila* was grown for 48 h in either mock or NanH1 pretreated GMM, as described above. Five μL of each culture was spread on a poly-L-lysine-coated coverslip and air dried for 20-30 min at RT. Bacteria were then fixed for 72 h with 1 mL of primary fixative: 2% (v/v) formaldehyde + 0,5% (v/v) Glutaraldehyde + 0.15% (w/v) Ruthenium Red in 0.1 M phosphate buffer pH 7.4. After two washes with 0.1 M phosphate buffer, pH 7.4, this was followed by a 2 h fixation in a post-fixative: 1% Osmium tetroxide + 1.5% (w/v) Ferocyanide in 0.065 M phosphate buffer pH 7.4. After a single wash with MilliQ, samples were dehydrated during consecutive 30 min steps with increasing concentrations of ethanol per step:50%, 70%, 80%, 95%, and twice with 100% ethanol, respectively. This was followed by 50% hexamethyldisilazane (HMDS) in ethanol and twice with100% HMDS. Following fixation and dehydration, samples were air dried overnight at RT and placed on EM thuds, coated with 6 nm gold and imaged using a FEI Scios FIB - Dual Beam SEM (ThermoFisher) at 5 kV.

### Bacterial length measurements

After acquisition of the fluorescent images, as described above, the length of 5 individual bacteria per field of view (FOV) was measured using Fiji software.^31^ For each biologically independent experimental replicate, 5 FOVs were assessed for a total of 75 bacteria per condition.

### Recombinant sialidase cloning and expression

#### Cloning of NanH1 in an expression vector

The NanH1 coding sequence (Allo_00594) of *Allobaculum mucolyticum* (Genbank accession number: JAHUZH010000000) lacking the predicted signal peptide (amino acids 1-29) was amplified from genomic DNA (isolated using the High Pure Template kit, Roche) by PCR using the Phusion Hot start II High Fidelity DNA Polymerase (Thermo Fisher Scientific) and primers NanH1-rec-F and NanH1-rec-R, both encoding a BsiWI restriction site (Table 1). The resulting PCR product was digested with BsiWI and ligated in frame with a N-terminal 3 x FLAG-tag and a C-terminal 6 × histidine tag into a modified and BsiWI-digested pET101/D-TOPO expression vector (Thermo Scientific), yielding pET101-NanH1.

#### Cloning of NanHIΔRIP

The NanH1 sequence was annotated using the Conserved Domains NCBI database (available at: www.ncbi.nlm.nih.gov/Structure/cdd/wrpsb.cgi). Seven putative catalytic amino acid residues within the sialidase domain likely form the active pocket and were identified based on similarity with other available sequences. Based on literature, the RIP motif (Arg449-Ile450-Pro451) was deemed part of a catalytic arginine triad important for the catalytic activity of the protein.^32^ Therefore, this site was selected for deletion. An inverse PCR was designed, using primers NanH1-RIP-rec-F NanH1-RIP-rec-R, with the forward primer inducing a 9-bp deletion in the DNA sequence encoding the RIP motif (CGAATCCCA). The inverse PCR product was amplified from pET101-NanH1, after which the template was removed by treatment 20 U of DpnI (NEB), a methylation-sensitive restriction enzyme. The DpnI-treated PCR products were further purified using agarose gel electrophoresis. Bands of the correct size were excised from the gel and purified using the GeneJET Gel Purification Kit (ThermoFisher Scientific), following manufacture’s protocol. The purified DNA fragment was blunt-end ligated using a T4 DNA ligase (ThermoFisher) resulting in plasmid pET101 – NanH1ΔRIP.

#### Cloning of NanH2

The NanH2 coding sequence (Allo_01388) was PCR amplified in similar fashion as NanH1, also lacking the predicted signal peptide (amino acids 1-29), using PCR primers NanH2-rec-F and NanH2-rec-R, both encoding a SmaI restriction site (Table 1). The resulting PCR product was digested with SmaI and ligated in frame with SmaI a N-terminal 3xFLAG-tag and a C-terminal 6xhistidine tag into a modified and SmaI-digested pET101/D-TOPO expression vector (Thermo Scientific), yielding pET101-NanH2.

All plasmids were checked for a correct orientation of the insert and sequences were confirmed by Sanger sequencing (Macrogen).

#### Protein expression and purification

The NanH1, NanH1ΔRIP and NanH2 expression vectors were transformed into *E. coli* BL21 (DE3) (Thermo Scientific) by heat-shock and plated on LB plates with 100 μg/mL ampicillin. Transformants were cultured in LB with 100 μg/mL ampicillin at 37°C and 160 RPM to an OD_600_ of 0.5 after which protein expression was induced by adding 1 mM IPTG (Thermo Scientific) for 4 h. Recombinant protein was purified using established lab protocols. In short, bacteria were pelleted (4,000 x *g*, 15 min, 4°C) and resuspended in 10 mL native buffer (50 mM Tris, 300 mM NaCl, pH8, 4°C). Lysozyme (200 μg/mL) and PMSF (1 mM) were added and mixtures were incubated for 15 min on ice. RNase and DNase (Roche, both at 5 μg/mL) were added and the mixture was incubated for another 10 min. Bacteria were subjected to three consecutive rounds of sonication, snap-freezing in liquid nitrogen and thawing (waterbath, 37°C). Sonication consisted of 3 × 10 s per round. Next, intact cells and cell debris were removed by centrifugation (4,000 x *g*, 30 min, 4°C) and imidazole was added (10 mM) to the supernatant. Fast protein liquid chromatography (FPLC) of this fraction was conducted using a 5 mL HiTrap Chelating HP Sepharose column on the ÄKTA FPLC protein purification system (GE Healthcare, Uppsala, Sweden). The proteins were eluted with 10 mL of starting buffer A (native buffer + 10 mM imidazole) followed by a gradient of buffer B (native buffer + 500 mM imidazole) at a flow rate of 5 mL/min, detected by absorbance at 280 nm, and collected in 1.8 mL fractions. Collected fractions were checked for purity using SDS-PAGE. Pure fractions were pooled and dialyzed overnight at 4°C against 5 L of PBS buffer. Glycerol was added at a final concentration of 10%, protein concentration was measured on Nanodrop, adjusted to 500 μg/mL and stored at −20°C until use.

**Table 1.**
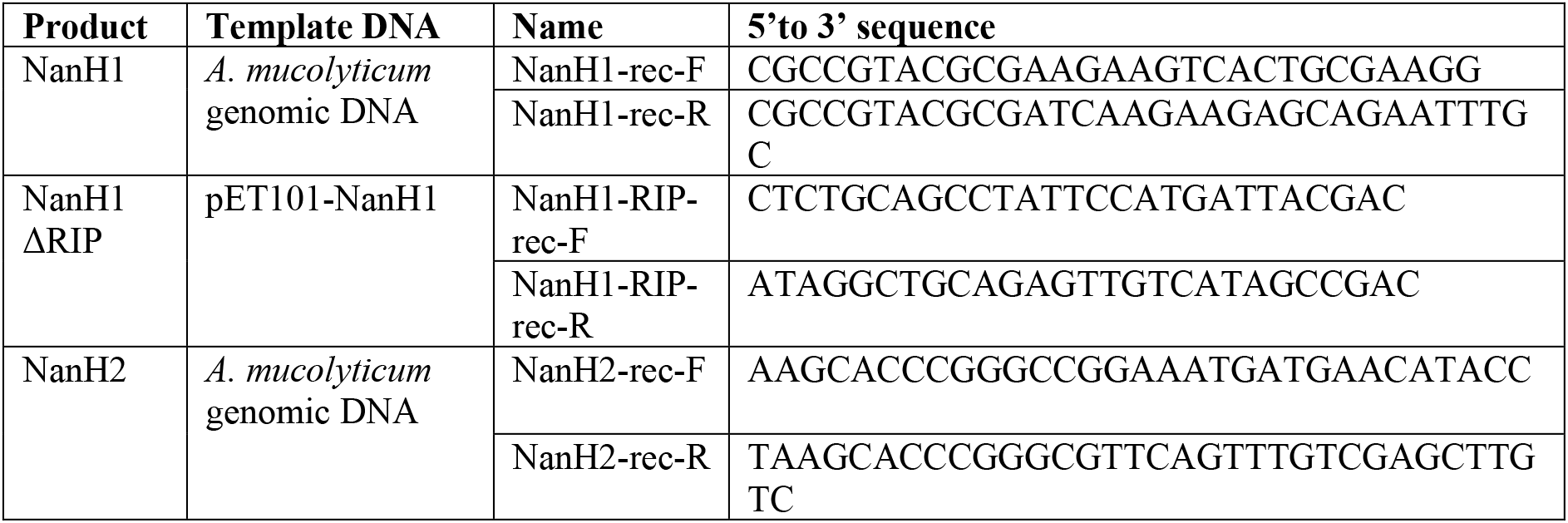
Primers used in this study

## Results

### Growth of *A. muciniphila* is inhibited by conditioned medium from *Allobaculum*

Direct competition with other intestinal bacterial species may contribute to the reduced abundance or absence of *A. muciniphila* in the intestinal tract. To identify inhibitory factors released by competing intestinal bacterial species, *A. muciniphila* cultures were supplemented with cell-free conditioned culture media from a panel of fourteen mucosal bacterial species isolated from patients with IBD that covered all major bacterial phyla (Figure 1A).

**Figure 1.**
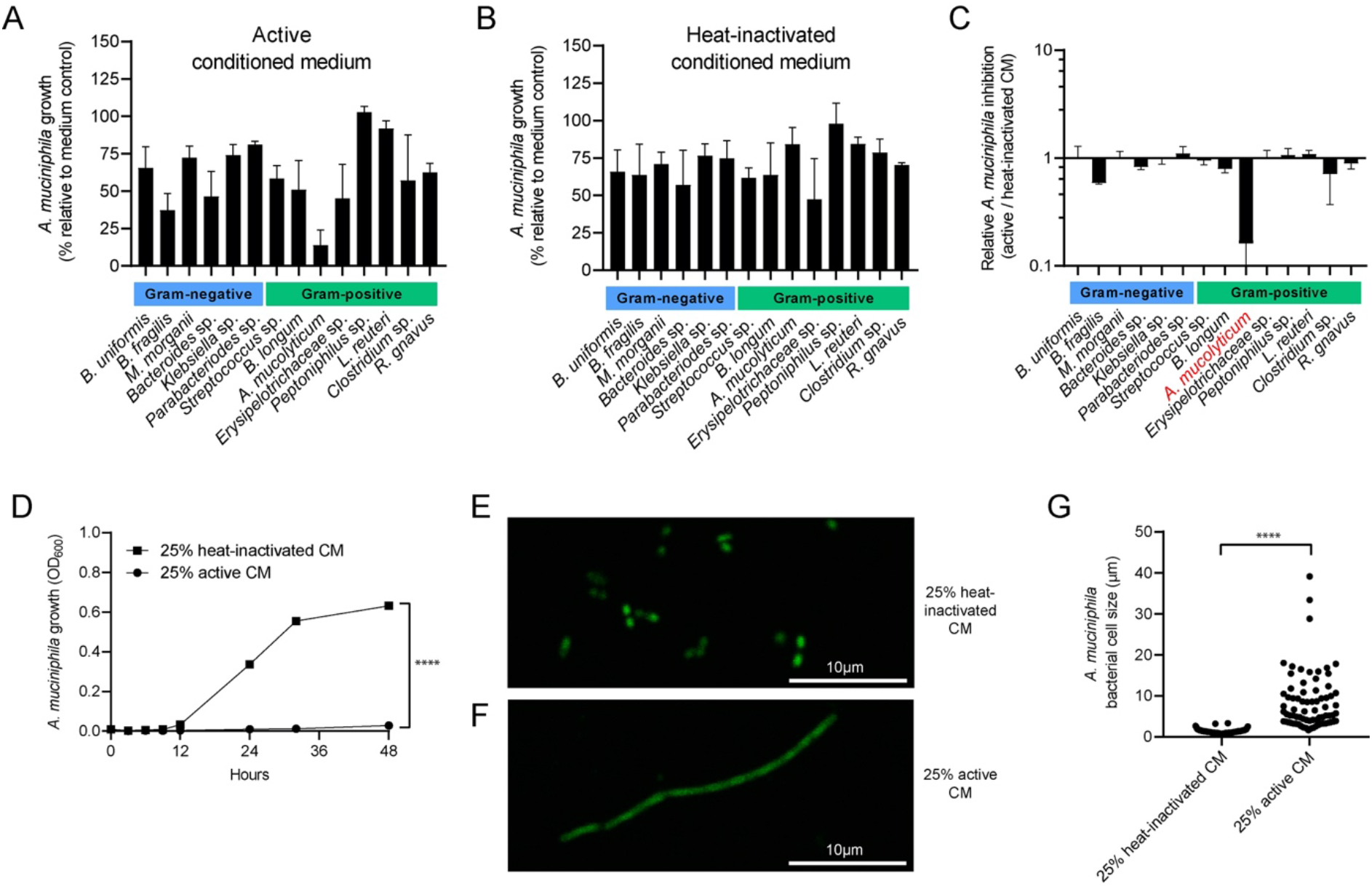
*A. muciniphila* is inhibited by *Allobaculum conditioned medium*. *A. muciniphila* growth was assessed after 48 h by measuring optical density at 600 nm (OD_600_). Bacteria were grown in Gut Microbiota Medium (GMM) supplemented with either active (A) or heat-inactivated (B) conditioned media (CM) from 14 different bacteria at a final concentration of 25%. The ratio between the final optical density values obtained with active or heat-inactivated media is displayed (C). (D) *A. muciniphila* growth curve showing growth over a 48 h period in the presence of active or heat-inactivated *Allobaculum* CM. Morphology of *A. muciniphila* was assessed after 48 h of growth in either heat-inactivated (E) or active (F) *Allobaculum* CM using confocal microscopy on syto-9 nuclear stained bacteria. Bars indicate 10 μm. Confocal images were used to measure bacterial cell length using ImageJ software (G). Per condition the length of 75 bacteria was recorded: five bacteria from five different fields of view per independent bacterial culture. Graphs depict the mean values ± SD acquired from three independent experiments. Statistical significance in D was determined for the 48 h time point by a *t* test. In G, statistical significance was determined using a Mann–Whitney U test. ****, *P* < 0.0001.

In order to differentiate between active growth inhibition and passive inhibition through selective nutrient depletion from the added conditioned culture media, a parallel set of *A. muciniphila* cultures were supplemented with heat-inactivated conditioned culture media from the fourteen bacterial species (Figure 1B). While the addition of heat-inactivated conditioned culture media from most bacterial species resulted in similar diminished growth of *A. muciniphila*, only one species, *Allobaculum mucolyticum* (for clarity, hereafter referred to as *Allobaculum*), showed strong inhibitory activity against *A. muciniphila* that was neutralized by heat-inactivation (Figure 1C). Closer inspection of the kinetics of the inhibition revealed that *A. muciniphila* did not enter the early-logarithmic growth phase in the presence of *Allobaculum* conditioned media, with minimal growth observed after culturing for 48 h. In the presence of heat-inactivated conditioned medium the growth of *A. muciniphila* was similar to growth in GMM control medium in which *A. muciniphila* entered a logarithmic growth phase after 9-12 h before reaching a maximum optically density after about 36-48 h (Figure 1D). *A. muciniphila* cultured in control GMM media or in media supplemented with heat-inactivated *Allobaculum* conditioned media displayed the typical rod-shaped morphology with an average bacteria lengths of 1.21 μm (Figure 1E,G). In contrast, *A. muciniphila* cultured in active *Allobaculum* conditioned media for 48 h displayed a severely elongated cell morphology with median bacterial length almost four times as long as control bacteria (5.73 μm), with some bacteria even reaching lengths of >30 μm (Figure 1F,G). Combined, these results suggests that *Allobaculum* releases a heat-sensitive inhibitory factor that severely hampers the growth of *A. muciniphila*.

### The inhibitory fraction of *Allobaculum-conditioned* media is enriched in glycolytic enzymes

Given the potent effects of *Allobaculum* conditioned medium on *A. muciniphila* growth and morphology, we set out to identify the inhibitory factor(s) secreted by this bacterium. To assess the approximate molecular size of the growth inhibitory factor(s) in the *Allobaculum* conditioned medium, size-fractionation using Amicon spin filters was performed. Testing of the different fractions for *A. muciniphila* growth inhibitory activity demonstrated potent inhibition for >50 kDa fractions (Figure 2B). As the size and heat-sensitivity of the inhibitory factor(s) may indicate a proteinaceous nature, we pre-treated the *Allobaculum* conditioned medium with proteinase K. This treatment rescued the growth of *A. muciniphila* (Figure 2C). Together, these results show that the growth of *A. muciniphila* is severely inhibited by one or more heat-sensitive proteins that are larger than 50 kDa.

**Figure 2.**
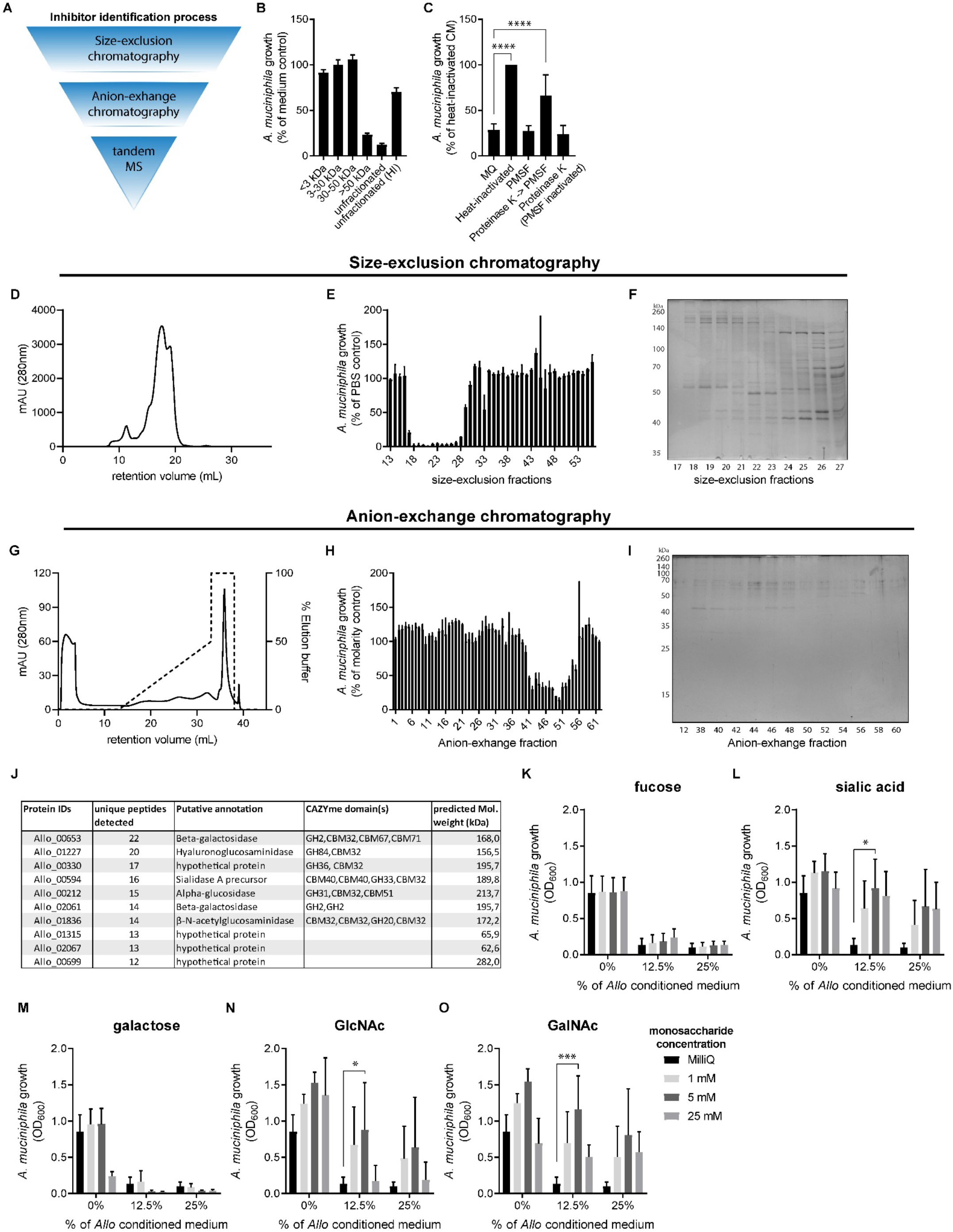
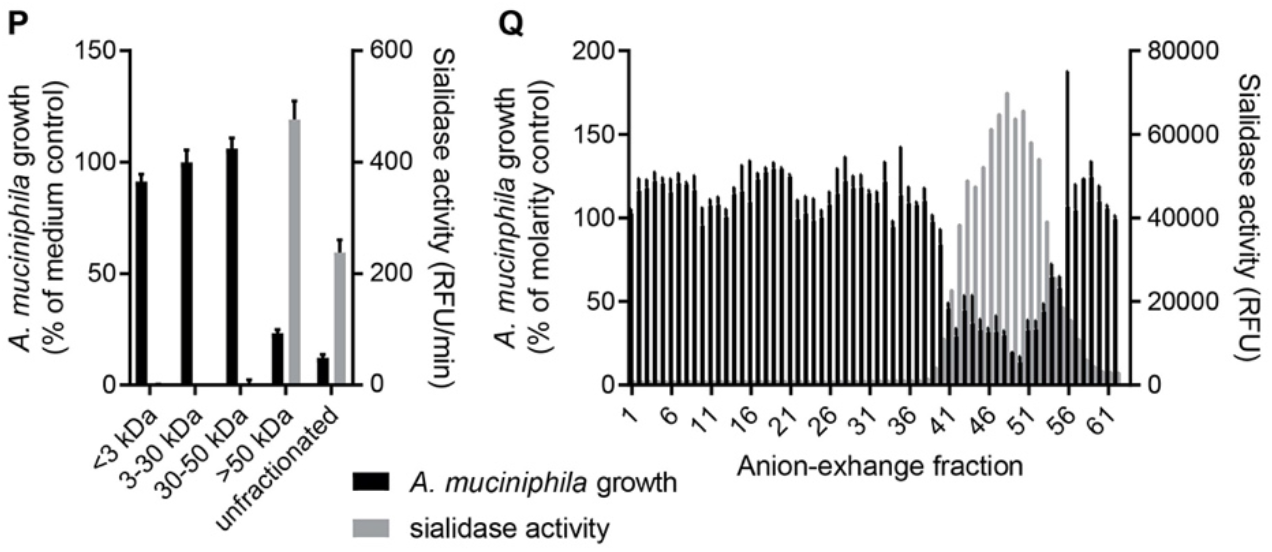
Identification of inhibitors in *Allobaculum* conditioned medium. (A) Overview of different steps in *Allobaculum* conditioned medium fractionation and inhibitor identification. (B) *A. muciniphila* growth was assessed after 48 h in the presence of 25% size-fractionated *Allobaculum* CM. Growth is depicted as a percentage of Gut Microbiota Medium (GMM) control. Size-fractionation was performed on Amicon spin-columns. (C) *A. muciniphila* growth (OD_600_) after 48 h in presence of 25%*Allobaculum* CM that was pretreated with Proteinase K. A 40x concentrated, >50 kDa fraction of *Allobaculum* CM was further fractionated using a size-exclusion column. (D) Chromatogram of size-exclusion fractionation showing retention volume (X-axis) and protein concentration as determined by measuring absorbance at 280 nm (Y-axis). (E) *A. muciniphila* growth at 48 h, as a percentage of PBS control, in the presence of the different size-exclusion fractions at a final concentration of 50%. Inhibiting fractions 17-27 were separated using SDS-PAGE and stained with a Coomassie stain (F). Inhibiting size-exclusion fractions 17 to 28 were pooled and separated further using anion-exchange chromatography. (G) Chromatogram of anion-exchange fractionation showing retention volume (X-axis) and protein concentration as determined by measuring absorbance at 280 nm (Y-axis). (H) *A. muciniphila* growth at 48 h, as a percentage of molarity control, in the presence of the different anion-exchange fractions at a final concentration of 25%. Inhibiting and non-inhibiting anion-exhange fractions were separated using SDS-PAGE and stained with a Coomassie stain (I). The strongest inhibiting anion-exhange fraction 50, and a non-inhibiting fraction 12 were analysed using LC-MS/MS and the detected peptides were aligned against the *Allobaculum* genome. The table (J) shows the ten most abundant proteins, as determined by the number of unique peptides per protein and their putative annotation based on automated annotations using Prokka and the dbCAN2 metaserver. *A. muciniphila* growth (OD_600_) after 48 h in presence of 0, 12.5 or 25% *Allobaculum* CM and 0, 1, 5 or 25 mM of either (K) D+fucose, (L), sialic acid (Neu5Ac), (M) D+galactose, (N) N-acetylglucosamine (GlcNAc) or (O) N-acetylgalactosamine (GalNAc). (P) The size-fractionated and (Q) anion-exchange fractions from a 48 h culture of *Allobaculum* grown in Gut Microbiota Medium, matching the fractions in figure 2B and 2H, respectively, were incubated with fluorescent 4-methylumbelliferone linked α-N-acetylneuraminic acid (sialic acid) for 30 minutes at 37°C. Fluorescence was recorded over time (P) or at the end-point (Q). All results are based on at least three experimentally independent replicates, each with two or three technical replicates per condition. Bar plots represent the mean ± SD. Statistical analysis was performed using GraphPad Prism, using a one-way (C) or two-way (K-O) ANOVA with Dunnett’s multiple comparison test. * = P < 0.05, ** = P < 0.01, *** = P < 0.001, **** = P < 0.0001.

In order to identify the inhibitory protein(s), a 40x concentrated >50 kDa *Allobaculum* supernatant fraction was subjected to high-resolution size-exclusion chromatography (Figure 2D). Separate fractions were collected and added to *A. muciniphila* cultures to a final volume of 50% after which *A. muciniphila* growth was assessed after 48 h. This experiment showed that fractions 17 to 28, corresponding to retention volumes between 7.5 and 13.6 mL, had the strongest inhibitory effect on *A. muciniphila* growth (Figure 2E). These fractions corresponded with a minor single peak in the chromatogram but still contained a variety of different proteins as shown by SDS-PAGE analysis (Figure 2F).

As a next step, the inhibiting size-exclusion fractions 17 to 28 were pooled and separated further using anion-exchange chromatography (Figure 2G). Again, fractions were collected and tested for *A. muciniphila* growth inhibition at a final concentration of 25% (v/v). As successive fractions contained increasing amounts of eluting buffer and therefore a higher concentration of NaCl, appropriate molarity medium controls were used to compare each fraction. This showed that anion-exchange fractions 41 to 55, corresponding to retention volumes between 27 and 34.5 mL, reduced growth of *A. muciniphila*, with fraction 50 causing the strongest growth reduction of more than 85% (Figure 2H). The inhibitory fractions corresponded with the third rather small peak in the anion-exchange chromatogram. SDS-PAGE analysis of the inhibitory fractions showed several bands (Figure 2I) but noticeably fewer than in Figure 2F, highlighting the effectiveness of the anion-exchange chromatography.

To learn more about the composition of the inhibitory fractions, the anion-exchange fraction with the strongest inhibitory activity (fraction 50) and a non-inhibiting fraction (fraction 12) were analysed using mass spectrometry. Detected peptides were aligned against the *Allobaculum* genome. The ten most abundant proteins detected in fraction 50, as determined by the number of unique peptides per protein, are displayed in Figure 2J. All these proteins had a molecular mass larger than 100 kDa. Automated annotation of the detected proteins suggested that three proteins were of unknown function, and a majority (7 out of 10) were putative glycosidases that contain one or multiple carbohydrate active enzyme (CAZY) domains. We have previously demonstrated that a number of these *Allobaculum* glycosidases are likely to be involved in mucin glycan degradation (Thesis Chapter 3).

### The secreted *Allobaculum* sialidase NanH1 inhibits the growth of *A. muciniphila*

To test whether glycosidases in the *Allobaculum* supernatant contributed to the inhibition of *A. muciniphila* growth, *A. muciniphila* was grown in the presence of 25% *Allobaculum* conditioned medium supplemented with one of five different monosaccharides often found in host-derived *O*-glycans: D+fucose, sialic acid (Neu5Ac), D+galactose, N-acetylglucosamine (GlcNAc) or N-acetylgalactosamine (GalNAc) (Figure 2K-O). We hypothesized that by adding these monosaccharides at high concentrations glycosidases present in the *Allobaculum* supernatant could be (partially) inhibited, leading to a rescue of *A. muciniphila* growth. In the presence of the highest concentration tested, D-(+)-fucose and D-(+)-galactose did not alleviate growth inhibition of *A. muciniphila* (Figure 2K & M). On the other hand, the presence of 5 mM Neu5Ac, GlcNAc or GalNAc rescued *A. muciniphila* growth (Figure 2L, N, O). Notably, when the concentration of either GlcNAc or GalNAc was increased to 25 mM, this again reduced the growth of *A. muciniphila*, suggesting potential toxicity/growth inhibitory effects of these substrates at higher concentrations. This effect was not seen with 25 mM of Neu5Ac. The consistent effect of sialic acid supplementation and the abundant presence of a sialidase in the inhibitory fraction as detected by MS (Figure 2J) led us to test for the presence of sialidase activity in the *Allobaculum* conditioned medium. Using the fluorescent 4-MU-NANA substrate, high levels of sialidase activity could indeed be detected in the inhibiting size-exclusion and anion-exchange fractions (Figure 2P & Q). Moreover, the levels of sialidase activity highly correlated with the level of inhibition of *A. muciniphila* growth. This combination of data suggests that the *Allobaculum* sialidase could play a role in the inhibition of *A. muciniphila* growth.

### Inhibition of *A. muciniphila* growth by the *Allobaculum* sialidase NanH1 and other bacterial sialidases

Inspection of the sequence of the sialidase gene (Allo_00594) of *Allobaculum* demonstrated that it encodes a 1,762 amino acid protein with six domains including a non-viral sialidase domain (Figure 3A). This sialidase is further referred to here as NanH1. The *Allobaculum* NanH1 protein and NanH1ΔRIP, a generated construct with a mutation in the active site, were expressed in *E. coli* after which sialidase activity was assessed using the 4-MU-NANA substrate (Figure 3B-C). Recombinant *Allobaculum* NanH1 demonstrated potent sialidase activity on this substrate, whereas this was completely absent after heat-inactivation or in NanH1ΔRIP. Moreover, the sialidase activity could be inhibited in a dose-dependent manner using 25 mM Neu5Ac or the sialic acid analogue oseltamivir phosphate (Figure 3C).

**Figure 3.**
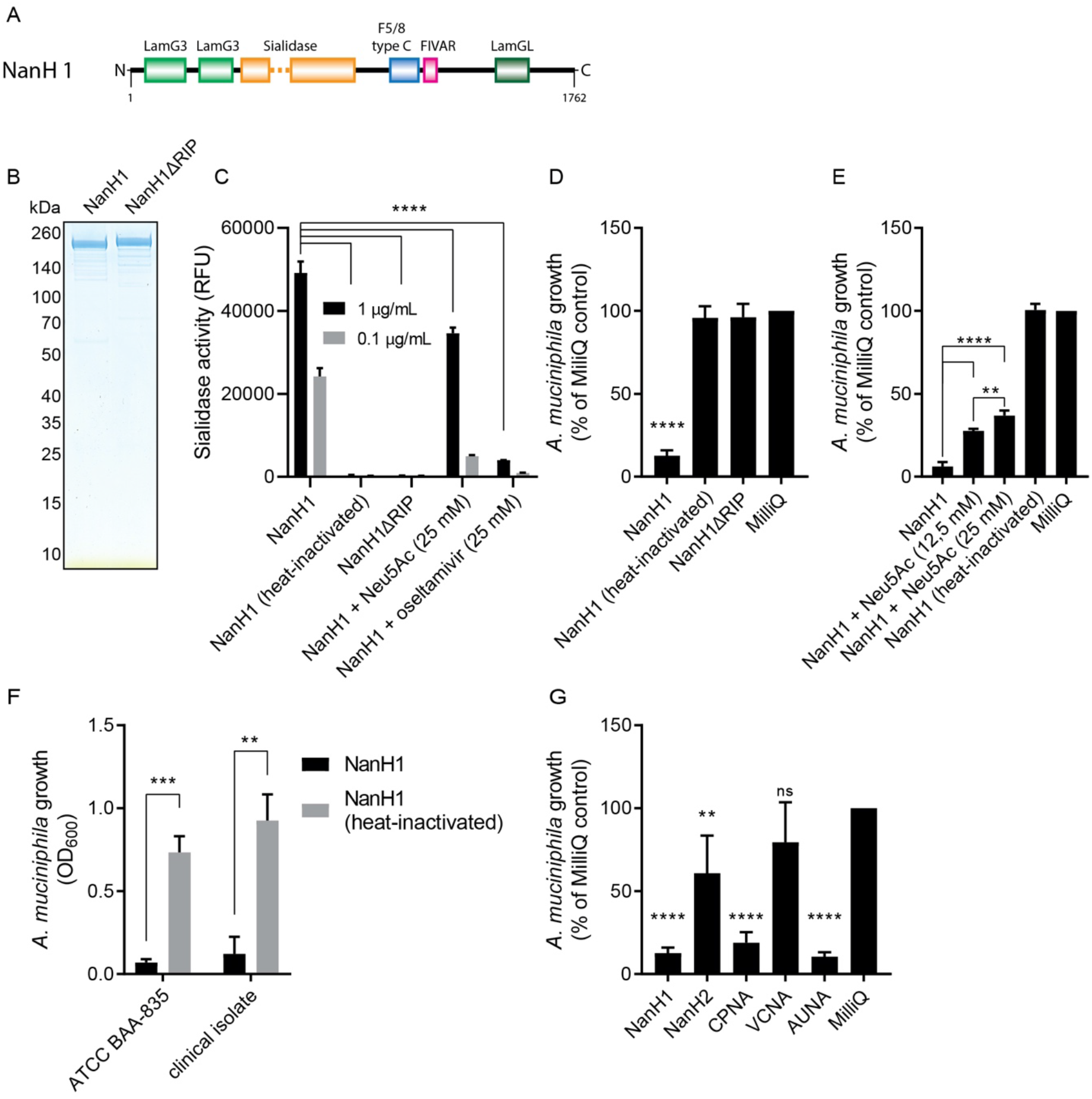
*Allobaculum* sialidase NanH1 and other bacterial sialidase inhibit the growth of *A. muciniphila*. (A) Domain architecture of NanH1. The displayed domains are those identified by BLAST search and are drawn to scale with polypeptide backbone. The number of amino acids in the backbone is indicated at the bottom. (B) Coomassie stain of recombinant NanH1 and NanH1ΔRIP separated by SDS-PAGE. (C) Sialidase activity of recombinant sialidases was determined by incubation with 4-methylumbelliferone linked α-N-acetylneuraminic acid (sialic acid) for 30 minutes at 37°C. Fluorescence was recorded over time. Figures D-G all indicate the growth of *A. muciniphila* after 48 h in GMM with following experimental conditions: (D) Growth as percentage of control in presence of recombinant NanH1 constructs (2.5 μg/mL). (E) Growth as percentage of control in presence of recombinant NanH1 (2.5 μg/mL) and sialic acid (Neu5Ac). (F) Growth (OD_600_) of *A. muciniphila* type strain (ATCC BAA-835) and clinical *A. muciniphila* isolate in the presence of active or heat-inactivated NanH1 (2.5 μg/mL). (G) Growth as percentage of control in presence of a panel of sialidases of which the enzyme activities were normalized to match the sialidase activity of NanH1 (2.5 μg/mL) on the 4-MU-NANA substrate. All results are based on at least three experimentally independent replicates, each with two or three technical replicates per condition. Bar plots represent the mean ± SD. Statistical analysis was performed using GraphPad Prism, using a two-way (C) or one-way (D, E, G) ANOVA with Tukey’s multiple comparison test. In figure F statistical significance was determined using and unpaired *t* test * = P < 0.05, ** = P < 0.01, *** = P < 0.001, **** = P < 0.0001.

Next, we checked whether the recombinant sialidase was capable and sufficient to inhibit the growth of *A. muciniphila*. Culturing *A. muciniphila* for 48 hours in media supplemented with recombinant NanH1 greatly reduced the growth of *A. muciniphila*, whereas heat-inactivated NanH1 or NanH1ΔRIP did not (Figure 3D). To confirm that the growth inhibition was caused by the sialidase activity rather than by one of the other domains present in NanH1, Neu5Ac was supplemented to the culture medium. This rescued the growth of *A. muciniphila* in a dose-dependent manner (Figure 3E), similar to what was seen for complete *Allobaculum* supernatant (Figure 2L). In order to test whether this sialidase-mediated inhibition is conserved across different strains of *A. muciniphila*, the type strain (ATCC BAA-835) was compared to a second, clinical *A. muciniphila* isolate. When recombinant NanH1 was added to a culture of this clinical isolate the growth of the isolate was also efficiently inhibited, suggesting that the mechanism underlying the growth inhibition is conserved among multiple strains. Together, these findings strongly suggests that the sialidase activity identified in the *Allobaculum* conditioned medium is due to NanH1 and that the sialidase activity is sufficient to inhibit *A. muciniphila* growth.

To assess the uniqueness of the inhibitory effect of the *Allobaculum* NanH1 on *A. muciniphila* growth, we tested four other bacterial sialidases for growth inhibitory activity: three commercially available bacterial sialidases from *Clostridium perfringens* (CPNA), *Vibro cholerae* (VCNA), and *Arthobacter ureafaciens* (AUNA) and NanH2, which originates from the same *Allobaculum* strain as NanH1. Notably, when added to *A. muciniphila* cultures at equal levels of enzyme activity, these sialidases demonstrated differential inhibitory activities. CPNA and AUNA inhibited the growth of *A. muciniphila* to a similar extend as *Allobaculum* NanH1, whereas *Allobaculum* NanH2 and VCNA were much less efficient inhibitors (Figure 3G). These results show that the inhibition of *A. muciniphila* is not restricted to the *Allobaculum* NanH1 sialidase and that differences in substrate specificity of sialidases likely determines the efficiency with which they can inhibit the growth of *A. muciniphila*.

### The *Allobaculum* sialidase NanH1 inhibits *A. muciniphila* growth through nutrient modification

Next, we investigated how sialidases inhibit the growth of *A. muciniphila*. While sialic acid may be present on the surface of *A. muciniphila*, the culture medium represents the largest source of sialylated substrates that could serve as target for the NanH1 sialidase. To examine whether nutrients within the culture medium were the prime target for NanH1 and its inhibiting effects on *A. muciniphila* growth, culture medium was pretreated for 16 h at 37 °C with NanH1 or heat-inactivated NanH1 as a control. After all remaining sialidase activity was eliminated by heat-inactivation, the treated culture medium was examined for its effect on *A. muciniphila* growth. Sialidase pretreatment of the culture medium inhibited the growth of *A. muciniphila* to the same extent as when active NanH1 was present during growth (Figure 4A). Additionally, both the growth kinetics and the morphological changes of *A. muciniphila* that were observed with *Allobaculum* conditioned medium, could be recapitulated by sialidase pretreatment of the culture medium (Figure 4A-C). In sialidase pretreated medium, the median length of *A. muciniphila* was 4.23 μm versus 1.31 μm in mock treated medium (Figure 4C). Together, these results indicate that *Allobaculum* NanH1 targets one or more components of the culture medium and that desialylation of this component results in *A. muciniphila* growth inhibition.

**Figure 4.**
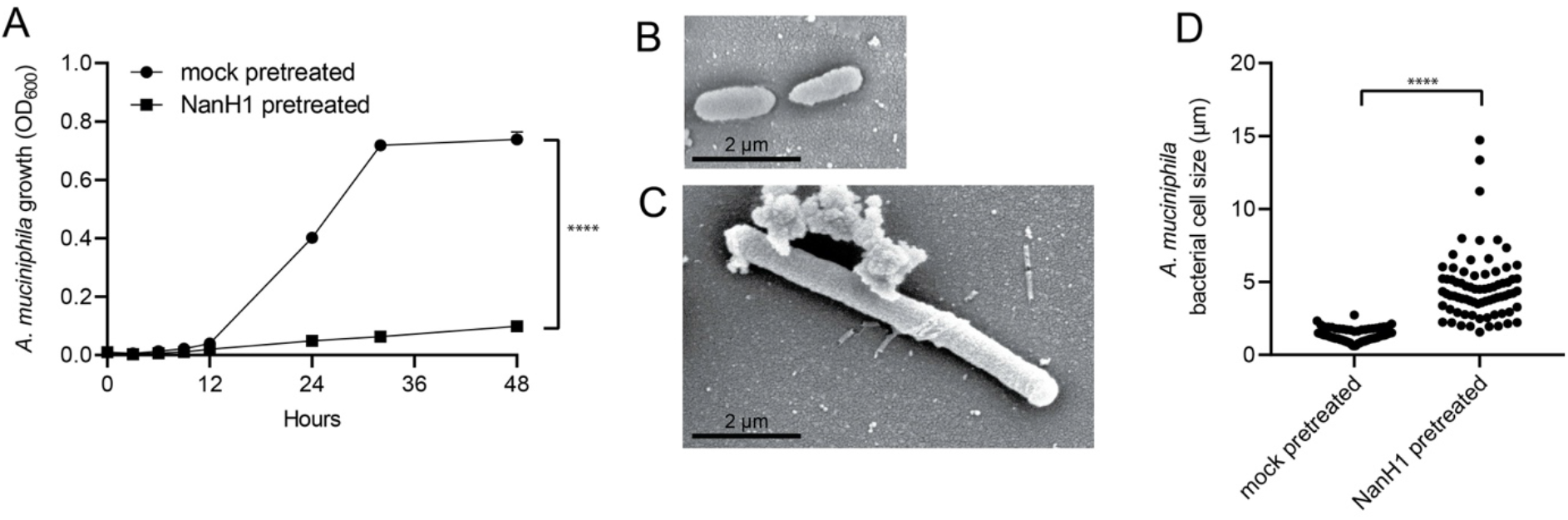
Growth of *A. muciniphila* is inhibited by pretreatment of growth medium with *Allobaculum* sialidase. (A) *A. muciniphila* growth curve over a 48 h period in the presence of NanH1 pretreated or mock pretreated Gut Microbiota Medium (GMM). Morphology of *A. muciniphila* was assessed after 48 h of growth in either (B) mock pretreated or (C) NanH1 pretreated GMM using scanning electron microscopy. Bars indicate 10 μm. *A. muciniphila* cell size was measured in confocal images of syto 9 nuclear stained bacteria grown for 48 h, using ImageJ software (D). Per condition the length of 75 bacteria was recorded: five bacteria from five different fields of view per independent bacterial culture. Graph depicts the mean values ± SD acquired from three independent experiments. Statistical significance in A was determined for the 48 h time point by a *t* test. In D, statistical significance was determined using a Mann–Whitney U test. ****, *P* < 0.0001.

### NanH1 sialidase inhibits *A. muciniphila* growth through desialylation of casein *O*-glycans

Gut microbiota medium contains many different components, but only three components that are predicted to contain sialylated glycans: trypticase peptone (a pancreatic digest of milk casein), yeast extract and meat extract. To first identify whether these and other components in the medium are required for *A. muciniphila* growth, different media were formulated, each lacking one component. This demonstrated that, in addition to a basal medium, glucose and trypticase peptone are essential medium components, in contrast to yeast extract and meat extract, which were dispensable for *A. muciniphila* growth (Figure 5A).

**Figure 5.**
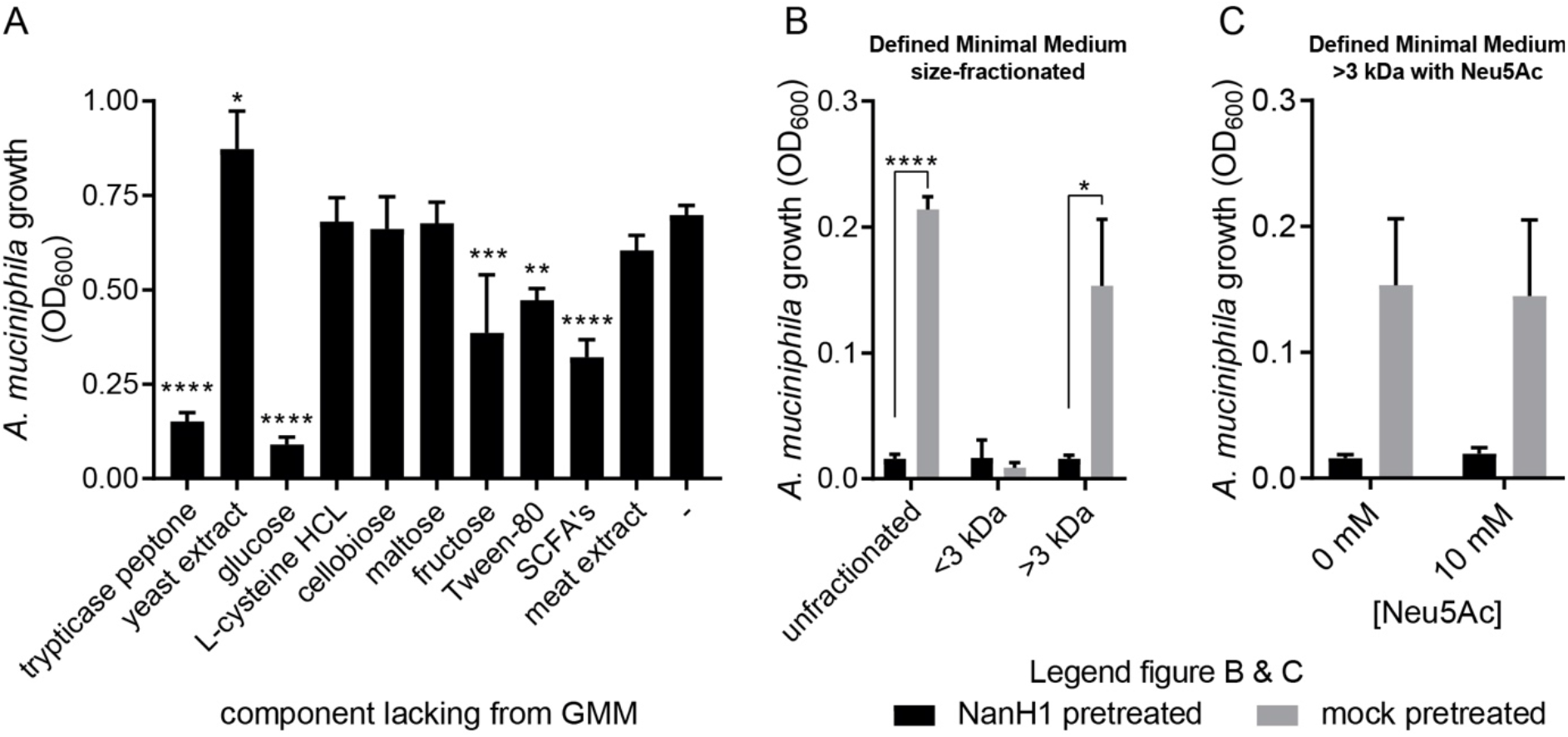
Identification of trypticase peptone as a target of *Allobaculum* sialidase and as an essential medium component for *A. muciniphila* growth. (A) *A. muciniphila* growth (OD_600_) after 48 h in variety of Gut Microbiota Medium formulations that lack one medium component at a time. (B) *A. muciniphila* growth (OD_600_) after 48 h in defined minimal medium (DMM) that was pretreated overnight with NanH1 (black bars) or a mock control (gray bars) and then size-fractionated. (C) *A. muciniphila* growth (OD_600_) after 48 h in NanH1 or mock pretreated >3 kDa DMM supplemented with 10 mM Neu5Ac. All results are based on at least three experimentally independent replicates, each with two or three technical replicates per condition. Bar plots represent the mean ± SD. Statistical analysis for figure A was performed using GraphPad Prism, using a one-way ANOVA with Dunnett’s multiple comparison test. Statistical significance in figure B was determined a *t* test. * = P < 0.05, ** = P < 0.01, *** = P < 0.001, **** = P < 0.0001.

A more defined version of the gut microbiota medium was formulated with a combination of trypticase peptone and glucose as the only major glycan and carbon sources. This defined minimal medium (DMM) was sufficient to support *A. muciniphila* growth, albeit to a lower maximum OD (Figure 5B). Pretreatment of DMM with NanH1 resulted in reduced growth of *A. muciniphila*, similar to what was seen with treatment of complete gut microbiota medium (Figure 5B). This indicates that the trypticase peptone is not only an essential nutrient source for *A. muciniphila* but also that the *Allobaculum* NanH1 sialidase likely targets sialylated glycans on casein, resulting in inhibition of *A. muciniphila* growth. To investigate whether the inhibitory effect is caused by a loss of sialic acid from trypticase peptone glycans or is the result of increased levels of free sialic acid, the NanH1-pretreated or mock-pretreated DMM was fractionated into <3 kDa and >3 kDa fractions. Multiple on-column wash steps ensured removal of any free sialic acid from the >3 kDa fractions. *A. muciniphila* was able to grow in either unfractionated or the >3 kDa fraction of untreated DMM, but not on NanH1-pretreated DMM or the <3 kDa fractions (Figure 5B). Moreover, when free sialic acid was supplemented to either the NanH1 or mock pretreated >3 kDa fraction, this did neither rescue nor inhibit the growth of *A. muciniphila* (Figure 5C). This demonstrates that the growth inhibition of *A. muciniphila* is not due to increased levels free sialic acid but rather due to the loss of sialic acid from casein glycans. Combined, these results demonstrate that *Allobaculum* inhibits *A. muciniphila* by secreting a sialidase that targets sialylated casein glycans that are critical for *A. muciniphila* growth.

## Discussion

Bacterial competition for nutrients and intermicrobial interactions are important determinants of the gut microbiota composition. In this study we set out to identify inhibitors of *A. muciniphila* growth that are secreted by bacterial competitors, as this may explain the reduced abundance of *A. muciniphila* in many diseases. Here we provide evidence that a sialidase from another intestinal mucin degrader, *Allobaculum*, can efficiently inhibit the growth of *A. muciniphila* through modification of the glycometabolic niche by targeting casein glycans that are required for growth.

We identified the sialidase-mediated inhibition of *A. muciniphila* by screening bacterial supernatants for inhibitory activity. The supernatant of *Allobaculum* clearly showed the most potent inhibition of *A. muciniphila* growth. Like *A. muciniphila*, *Allobaculum* secretes many glycosidases that can degrade (mucin) *O*-glycans (Chapter 3), showing that the metabolic programs of both bacteria are optimized for feeding on host glycans. This is important as it places these two anaerobic enteric bacteria in the same mucosal niche, where they would likely compete for space and nutrients. When competing for the same nutrients it can be imagined that bacteria evolve mechanisms to inhibit their competitors. However, as both *A. muciniphila* and *Allobaculum* are capable of producing sialidases, and sialidases are not known to act as inhibitors of bacterial growth, we did not anticipate to identify the *Allobaculum* NanH1 sialidase as the inhibitor of *A. muciniphila* growth^33^.

The NanH1 sialidase was identified as an inhibitor through a reductionist approach using multiple, consecutive fractionation methods. However, due to the sensitivity of mass spectrometric analyses this still left multiple candidate proteins, of which the majority were glycosidases that could be involved in mucin degradation. Due to the large number of candidate proteins we opted for an approach using the enzymes’ putative cognate ligands, the mucin monosaccharides, to act as competitive inhibitors. This proved successful in identifying potential candidates as we were able to rescue the growth of *A. muciniphila* using Neu5Ac, GlcNAc and GalNAc. This suggested that the glycosidases that target these monosaccharides might be involved in the inhibition of *A. muciniphila*. Nonetheless, these findings could also point to the importance of GlcNAc and GalNAc in *A. muciniphila’s* metabolism, as exogeneous supplementation of these metabolites was reported to be essential for *A. muciniphila* growth^34^. As multiple putative glucosaminidases were on our candidate list and, in this setup, it was difficult to delineate the actions of these enzymes from the effects of the monosaccharides, we instead focused on the sialidase as the addition of Neu5Ac also rescued growth, but Neu5Ac cannot be metabolized by *A. muciniphila*^35^.

Multiple lines of evidence indicate that the NanH1 sialidase is able, and by itself sufficient to inhibit the growth of *A. muciniphila*. As a first step, using a sialidase activity reporter, we showed a near perfect inverse correlation between the levels of sialidase activity in the inhibitory fractions of the *Allobaculum* conditioned medium and the growth of *A. muciniphila*. Second, and most convincingly, we showed that recombinant NanH1 was capable of inhibiting *A. muciniphila* growth, but not when its sialidase activity was inhibited by Neu5Ac or inactivated through mutation of the catalytic site or by heat. As an additional confirmation of the central role of sialidase activity in the inhibitory process we showed that several other bacterial sialidases can also act as inhibitors of *A. muciniphila* growth. Bacterial sialidases often function as virulence factors and are known to play important roles in the degradation of mucin glycans. However, according to our knowledge, this is the first description of a sialidase that acts as an inhibitor of bacterial growth.

As a next step we identified the target of this sialidase. We demonstrated that pretreatment of the medium with NanH1 was sufficient to inhibit the growth of *A. muciniphila*. This indicated that the inhibition is mediated through modifications of sialoglycans in the growth medium rather than through direct effects on the cell surface of *A. muciniphila*. Then, using different growth media formulations, we showed that trypticase peptone was required for *A. muciniphila* growth and, importantly, that desialylation of this substrate was sufficient to inhibit the growth of this bacterium. Trypticase peptone is a pancreatic digest of bovine *k*-casein. Casein is a major glycoprotein in both bovine and human milk, and after digestion of *k*-casein by gastric and pancreatic proteases the glycosylated C-terminal part of this protein is also referred to as glycomacropeptide (GMP)^37^. GMP contains multiple *O*-glycans, which are made up of terminally located sialic acid residues and underlying residues of galactose, GlcNAc and GalNAc. GlcNAc and/or GalNAc, as well as L-threonine, which is abundantly present in the polypeptide backbone, are essential nutrients required for *A. muciniphila* growth^34^. The fact that GMP contains this tailor-made combination of essential metabolites likely explains the requirement for the substrate in the GMM growth medium and may provide clues about the mechanism underlying the sialidase-mediated growth inhibition of *A. muciniphila*.

Several scenarios can be envisioned as to how the sialidase may cause the inhibition of *A. muciniphila* growth on GMP. A first explanation could be that presence of sialic acid on the *O*-glycans is required for efficient hydrolysis of the glycan or the peptide backbone. However, this conflicts with the consensus view that the rate of glycoproteolysis is generally increased after the removal of terminal sialic acids^38–40^. An example of this is the much higher enzyme activity of the *A. muciniphila O*-glycopeptidase, OgpA, towards non-sialylated substrates^41^.

A second explanation could be that glycan-associated but not free sialic acid provides essential cues for the utilization of the casein GMP substrate, or that high concentrations of free sialic acid lead to catabolic repression. Transcriptional repression of the enzymes required for the sequential degradation of the underlying glycoprotein would then prevent further growth. However, catabolic repression by free sialic acid is not likely, as we were able to rescue growth of *A. muciniphila* in the presence of active *Allobaculum* conditioned medium by adding an excess of free sialic acid (Figure 2L). Moreover, *A. muciniphila* was also not able to grow on NanH1 pretreated DMM from which the liberated sialic acids were removed (Figure 5B). Although we cannot rule out the possibility that only complete, sialylated *O*-glycans may provide essential metabolic cues for *A. muciniphila*, it is difficult to envision how this would allow the bacterium to acquire essential nutrients, like GlcNAc and L-threonine, in situations when it actively secretes its own sialidases^33^. Although we have not verified the contribution of *A. muciniphila’s* own sialidases to this process, they are likely required for the degradation and growth on the fully sialylated GMP.

A third explanation for the growth inhibition could be that *A. muciniphila*, grown in the presence of fully sialylated GMP, tightly regulates its metabolic programme to match the rate at which monosaccharides are released from sialylated glycans, and that this rate is determined by *A. muciniphila’*s own sialidases. A suddenly increased rate of desiaylation by an external sialidase, such as the *Allobaculum* sialidase, might disrupt this delicate balance and could lead to an increased rate of hydrolysis of the underlying glycan and a concomitant sudden increase in free *O*-glycan monosaccharides, such as galactose, GlcNAc and GalNAc. As the metabolic pathways for these substrates are highly intertwined, a sudden change in their availability could lead to metabolic dysregulation and a cessation of growth^35,42^. An example of this could be the toxic accumulation of sugar-phosphates, such as galactose-1-phosphate, which are linked to growth defects in wide variety of species across multiple kingdoms of life^43–45^. Van der Ark *et al*. have also demonstrated that these metabolic pathways are intricately linked to processes of bacterial growth and cell morphology, as *A. muciniphila* requires exogenous GlcNAc or GalNAc for peptidoglycan synthesis^34,42^. Similar to our observation in sialidase-pretreated medium, this study also observed a highly elongated morphology of *A. muciniphila* when it was grown in synthetic PT medium^42^. The authors suggested that it could be caused by an imbalance between the availability of glucose and GlcNAc in this medium. The fact that these different growth media induce a similar elongated phenotype of *A. muciniphila* may point towards a similar underlying mechanism of metabolic dysregulation.

Besides valuable new insights into the intricate metabolism and cell biology of *A. muciniphila*, our findings also shed a new light on the role of sialidases in microbial ecology. First of all, we demonstrate that GMP supports the growth of *A. muciniphila*. The fact that this highly abundant milk protein supports growth may not be entirely surprising given the fact that the *O*-glycans present on GMP are similar to mucin O-glycans. Furthermore, it also aligns with a recent report by Kostopoulos *et al*.^46^ who reported that *A. muciniphila* can also use human milk oligosaccharides for growth. Moreover, the fact that the highly abundant milk protein supports the growth of *A. muciniphila* can have important implications for the establishment of the early life microbiota. This notion is supported by the fact that GMP glycans also promote the growth of *Bifidobacterium longum* subsp*. infantis*, which colonizes the infant early after birth^47^. Many of the first bacterial colonizers in early life, including many of the *Bacteroides* and *Bifidobacteria* are known sialidase producers. Even though we did not see a significant inhibitory effect of these species on *A. muciniphila’s* growth *in vitro* (Figure 1A-C), sialidases from these bacteria or from pathobionts similar to *Allobaculum* could be important actors *in vivo* and affect *A. muciniphila’s* ability to establish itself within the neonatal intestinal ecosystem. Besides the effects on bacterial ecology, desialylation of GMP may also affect some of GMP’s other biological functions. Sialic acid residues on GMP were previously shown to neutralize enteropathogens as well as inhibit the LPS-induced proliferation of spleen lymphocytes^48–50^. Sialidase-mediated removal of these sialic acids could therefore impact intestinal homeostasis.

In addition to demonstrating the inhibitory effect of the *Allobaculum* NanH1 sialidase, we also demonstrated that this effect can be exerted by sialidases from other bacteria, such as *C. perfringens*.This implies that the findings from *Allobaculum* could potentially also be extended to other sialidase-producing bacteria living in the intestinal mucosal niche. This hypothesis is in line with findings by Png *et al.*^2^ who reported inverse correlations between the abundance of *A. muciniphila* and the sialidase-producing mucin degraders *Ruminococcus gnavus* and *Ruminococcus torques*, which are known to occupy the same niche as *A. muciniphila*. We did not detect an inhibitory effect of the cell free supernatant of *R. gnavus* (Figure 1) but this might be explained by strain-dependent differences in the absence or presence of sialidase genes, or an inability of *R. gnavus* to sufficiently express the sialidases in the culture medium^39^.

The fact that multiple isolates of *A. muciniphila* seem to be susceptible to sialidase-mediated growth inhibition suggests that the mechanisms underlying this susceptibility are conserved and that this trait may be essential or beneficial *in vivo*. A reduced growth rate can have major negative consequences for bacterial fitness in complex multi-species bacterial communities, where competition for space and nutrients serves as a strong selective pressure. As such the question may arise why this trait is evolutionarily conserved among multiple strains of *A. muciniphila*, even though it makes them vulnerable to sialidase-mediated growth inhibition^51^. One possible explanation is that *A. muciniphila* does not commonly encounter “external” inhibitory sialidases. However, this is unlikely given the abundance of sialidase-encoding bacteria in the mucosal niche, even when considering not all sialidases may have the required specificity. Another explanation could be that *A. muciniphila* is incapable of adapting to sialidase inhibition, as the mutations or altered gene expression required to adapt interfere with processes that are essential for cell division and survival. The dependency on exogenous GlcNAc/GalNAc for peptidoglycan synthesis may support this notion. Alternatively, it can be imagined that successful adaptation to sialidase-mediated inhibition may alter the bacterium’s physiology (e.g., capsule or membrane structure) in such a way that this renders it more susceptible to other even stronger selective pressures such as phage predation or host immune responses^52^.

The question remains whether our findings also apply in a more complex *in vivo* setting. This will depend on whether *A. muciniphila* and *Allobaculum*, or other sialidase-producing bacteria do indeed encounter each other in the same (mucosal) niche, and whether the sialidase is produced and able to exert the inhibitory influence under these conditions. The fact that both *A. muciniphila* and *Allobaculum* have the ability to degrade mucins and the fact they are both found to be highly IgA coated suggests that they may inhabit the same niche *in vivo*. Regardless, our bottom-up *in vitro* approach allowed us to identify and validate causal relationships between the growth of *A. muciniphila* and secreted products from other bacteria. This powerful approach can also be applied to any bacteria for which conditioned culture media are available. Moreover, the simplicity and efficacy of our experimental setup could allow for easy upscaling to a much larger number of bacterial supernatants, especially when combined with high-throughput culturomics techniques. Such a high-throughput screen will likely result in the identification of more growth inhibiting or growth promoting factors for *A. muciniphila* and other bacteria. As we have shown here, this approach can unveil hitherto unknown molecular mechanisms and provide important new insights in microbial cell biology and microbial ecology. These new insights may pave the way for new therapeutic approaches that prevent the loss of *A. muciniphila*.

