## Supplemental Table 1 for "Growth inhibition of *Akkermansia muciniphila* by a secreted pathobiont sialidase"

**Supplementary Table 1. Recipe for enriched Gut Microbiota Medium (GMM)**

| Component | Amount (500 mL) | Concentration | Comments | Vendor |
|---|---|---|---|---|
| Tryptone Peptone | 5 g | 10 g/L | | BBL Trypticase Peptone BD (211921) |
| Yeast Extract | 2.5 g | 5 g/L | | Bacto Yeast Extract BD (212750) |
| D-(+)-glucose - absent from basal medium | 1 g | 11 mM | | Sigma (G8270-1KG) |
| L-cysteine HCL | 0.5 g | 6.4 mM | | Sigma (C-1276) |
| D-(+)-Cellobiose - absent from basal medium | 0.5 g | 2.9 mM | | Sigma |
| D-(+)-Maltose - absent from basal medium | 0.5 g | 2.8 mM | | Sigma |
| D-(-)-Fructose - absent from basal medium | 0.5 g | 2.2 mM | | Sigma |
| Meat Extract | 2.5 g | 5 g/L | | Sigma |
| Phosphate buffer (see below) | 100 mL | 100 mM | 1 M stock solution pH 7.2 | |
| TYG salts solution (see below) | | | | |
| $CaCl_2$ | 0.5 mL | 0.8% | 8 mg/mL stock solution | Sigma |
| Vitamin K (menadione) | 0.5 mL | 5.8 mM | 1 mg/mL stock solution | Sigma |
| $FeSO_4 \cdot 7H_2O$ | 0.5 mL | 1.44 mM | 0.4 mg $FeSO_4 \cdot 7H_2O$/mL stock solution | Sigma |
| Histidine Hematin Solution (see below) | 0.5 mL | 0.1% | 1.2 mg hematin/mL in 0.2 M histidine solution | Sigma |
| Tween 80 | 1 mL | 0.05% | 25% stock solution | Sigma |
| ATCC Vitamin Supplement | 5 mL | 1% | | ATCC |
| ATCC Trace Mineral Supplement | 5 mL | 1% | | ATCC |
| Short-chain fatty acid supplement (see below) | 3.75 mL | | | |
| Resazurin | 2 mL | 4 mM | 0.25 mg/mL stock solution | Sigma |

| **Potassium Phosphate buffer recipe (1 M, pH 7.2)** | |
|---|---|
| 1 M $KH_2PO_4$ (68.045 g/500 mL) | 430 mL |
| 1 M $K_2HPO_4$ (174.18 g/L) | 1 L |

*add monobasic to dibasic to achieve pH 7.2, autoclave, store at room temperature*

| **TYG salts solution recipe** | **Amount (1 L)** | |
|---|---|---|
| $MgSO_4 \cdot 7H_2O$ | 0.5 g | Sigma |
| $NaHCO_3$ | 10 g | Sigma |
| NaCl | 2 g | Sigma |

| **Histidine solution recipe (0.2 M, pH 8)** | | |
|---|---|---|
| histidine | 2.1 g | Sigma |
| MilliQ | bring to 40 mL | |

*Set pH to 8.0 with 5 M NaOH, then bring to 40 mL*

| **Histidine-Hematin recipe (0.2 M)** | | |
|---|---|---|
| hematin | 24 mg | Sigma |
| histidin solution (0.2 M, pH 8) - see above | 20 mL | |

*Cover in aluminum foil, rotate overnight at 4°C. Filter sterilize, aliquot in 0.5 mL, store at -20°C*

| **Short-chain fatty acid supplement recipe** | **Amount (50 mL)** | |
|---|---|---|
| Acetic acid | 11.33 mL | |
| Sodium propionate Na salt | 5.21 g | Sigma |
| Sodium butyrate Na salt | 3.21 g | Sigma |
| MilliQ | bring to 50 mL | |
